# Distinct excitability of thalamocortical neurons correlates with the presence of cerebellar afferents

**DOI:** 10.1101/2023.05.26.542536

**Authors:** Myriam Moreno, Crystal Minjarez, Slobodan M. Todorovic, Nidia Quillinan

**Affiliations:** Department of Anesthesiology, 12801 E. 17th Ave. MS8130, Research 1 South, Aurora, CO 80045, USA; Neuronal Injury and Plasticity Program, 12801 E. 17th Ave. MS8130, Research 1 South, Aurora, CO 80045, USA

**Keywords:** intrinsic excitability, ventrolateral thalamus, cerebellar afferents, tonic firing, rebound firing

## Abstract

Thalamocortical (TC) neurons within the ventrolateral thalamus (VL) receive projections from the cerebellum and the basal ganglia (BG) to facilitate motor and non-motor functions. Tonic and rebound firing patterns in response to excitatory cerebellar and inhibitory BG inputs, respectively, are a canonical feature of TC neurons and plays a key role in signal processing. The intrinsic excitability of TC neurons has a strong influence on how they respond to synaptic inputs, however, it is unknown whether their afferents influence their firing properties. Understanding the input-specific firing patterns could shed light into movement disorders with cerebellar or BG involvement. Here, we used whole-cell electrophysiology in brain slices from C57BL/6 mice to investigate the firing of TC neurons with optogenetic confirmation of cerebellar or BG afferents. TC neurons with cerebellar afferents exhibited higher tonic and rebound firing rates than those with BG afferents. This increased firing was associated with faster action potential depolarization kinetics and a smaller afterhyperpolarization potential. We also found differences in the passive membrane properties and sag currents during hyperpolarization. Despite higher rebound firing in TC neurons with cerebellar afferents, there were no differences in T-type calcium channel function compared to those with BG inputs. These data suggest input-specific differences in sodium and SK, but not T-type calcium channels, impact firing properties in TC populations. Altogether, we showed that the pronounced divergence observed in TC neuron firing properties correlate with its heterogeneous anatomical connectivity, which could signify a distinct signal integration and processing by these neurons.

**Keypoints:**

- Thalamocortical neurons in the VL with cerebellar afferents have higher intrinsic tonic and rebound firing properties than those with basal ganglia afferents.
- Membrane resistance and action potential depolarization slope were different based on the presence of cerebellar afferents.
- Despite elevated rebound burst firing, T-type mediated currents did not correlate with increased firing in neurons with cerebellar afferents.

## Introduction

The ventrolateral (VL) thalamus receives, integrates and relays signals from the cerebellum and the basal ganglia to the motor cortex (Buford *et al*., 1996; Sommer, 2003; Kasten & Anderson, 2015; Hintzen *et al*., 2018). Thalamocortical (TC) neurons within the VL thalamus receive excitatory from the deep cerebellar nuclei and inhibitory input from the nucleus reticularis thalami (nRT), the internal segment of the globus pallidus (GPi), and substantia nigra (SNr) (Paz *et al*., 2010; Garcia-Munoz & Arbuthnott, 2015; Caligiore *et al*., 2017; Kim *et al*., 2017; Pelzer *et al*., 2017; Hintzen *et al*., 2018; Habas *et al*., 2019). Synchronous integration of excitatory and inhibitory input from the cerebellum and the basal ganglia by the thalamus allows for fine-tuned communication with the cortex. This circuit mediates complex and adaptive behaviors, including sensory motor control and optimization of motor sequences, decision-making, habit formation and implicit learning (Garcia-Munoz & Arbuthnott, 2015; Pelzer *et al*., 2017; Bostan & Strick, 2018).

The thalamus, cortex, cerebellum, and the basal ganglia (GPi, SNr, and striatum) are heavily interconnected (Pelzer *et al*., 2017; Hintzen *et al*., 2018) making it a complex circuitry to dissect when it comes to identifying molecular and cellular targets associated with motor-related disorders. A disruption due to injury or disease at any level of the circuitry can affect the output of the network. For instance, damage to the cerebellum affects the entire cerebello-thalamo-cortical pathway and causes motor coordination disorders such as ataxia and dystonia, as well as cognitive dysfunction (Schmahmann, 1996; Schmahmann *et al*., 2009; Stoodley *et al*., 2016; Stoodley & Schmahmann, 2018). Loss of dopaminergic neurons in the SNr leads to an altered inhibitory output to the VL thalamus and the striatum, affecting the basal ganglio-thalamic pathway, causing movement disorders characteristic of Parkinson’s disease and dystonia (Poewe *et al*., 2017; Shakkottai *et al*., 2017; Bove & Travagli, 2019; Qian *et al*., 2020).

The thalamic network regulates sensory transmission via fluctuations between tonic and rebound or burst firing modes (Sherman, 2001; Sorokin *et al*., 2017; Zeldenrust *et al*., 2018). Tonic firing is a regular mode of continuous spiking of action potentials throughout a depolarizing stimulus, whereas rebound burst firing consists of a brief cluster of action potentials triggered after a hyperpolarized state. Tonic firing is primarily mediated by a combination of sodium (Na^+^) and potassium (K^+^) channels (Rzhepetskyy *et al*., 2016; Stamenic & Todorovic, 2018). While rebound firing is primarily mediated by the activation of low-voltage-gated T-type Ca^2+^ and HCN channels following a hyperpolarization or GABAergic inhibitory which relieves them from inactivation (Kasten & Anderson, 2015; Leresche & Lambert, 2017). Ion-channel associated changes in the firing properties of thalamic neurons are implicated in a number of disease states. Studies in rodents have shown that injury to cortico-thalamic or spino-thalamic tracts leads to an increase in excitability of thalamocortical neurons (Wang & Thompson, 2008; Paz *et al*., 2010; Nielsen & Jensen, 2017). However, we lack a clear understanding of heterogeneity in intrinsic excitability within the VL and whether afferent projections may inherently influence tonic or rebound firing states that are typical of TC neurons.

Identification of differences in biophysical properties and excitability of thalamocortical neurons at a molecular level (ion-channels), could be pivotal towards understanding the mechanisms by which injury or disease alter motor function. Evidence of anatomic segregation of cerebellar and basal ganglia afferents onto thalamocortical neurons, along with gene expression gradients found at the single cell level within the VL (Roy *et al*., 2022), could suggest a strong association between specific thalamic properties and thalamic input. To better understand the physiology of these neurons in accordance with their afferents, we used whole-cell patch-clamp electrophysiology in *ex vivo* acute brain slices to investigate the intrinsic excitability of thalamocortical neurons with cerebellar or basal ganglia afferents. Our results revealed distinct firing rates and biophysical properties that are associated with either type of afferent. Furthermore, we identified various ion channels to determine their role in the observed excitability differences.

## Materials and Methods

### Animals

All experimental protocols were approved by the Institutional Animal Care and Use Committee (IACUC) at the University of Colorado and adhered to the National Institute of Health guidelines for the care and use of animals in research. Male and female, 8-12 weeks old, C57Bl/6N-Elite mice were purchased from Charles River Laboratories (CRL). All mice were permitted access to water and standard lab chow ad libitum with a 14/10-h light/dark cycles. For current clamp experiments in Figures 2 and 4(C-H), a total of 31 animals were used. For voltage clamp experiments in Figure 4 (inactivation and recovery from inactivation protocols), a total of 5 animals were used.

### Intracranial injections

Stereotaxic intracranial injections with AAV-hSyn-ChR2-mCherry (Addgene, 26976) or AAV-hSyn-CHIEF-mRuby (courtesy of the Aoto Lab, University of Colorado Denver Anschutz Medical Campus) were performed at least 3 weeks prior to electrophysiological experiments in 8-12 weeks old mice with a Nanoject II. The cerebellar nuclei were targeted by using the left superior cerebellar artery and superior cerebellar vein as a reference (Moreno et. al, 2022): x=-0.60-0.7, zeroed at the artery; y=0.5-0.6, zeroed at the vein; z=2.1 at a 31° angle). Virus was injected at a rate of 20 nL per pulse every 10 seconds for a total volume ∼80 μL of virus in the lateral/interposed nuclei and ∼100nL for SNr. Coordinates for SNr injections were based on bregma (x=1.5, y=3.3 and z=4.44). Injection site into SNr and Cb was confirmed for each animal by localizing mRuby fluorescence in the ipsilateral SNr or contralateral cerebellar nuclei, relative to right thalamus where recordings were performed.

### In vivo slice preparation

Mice were anesthetized with 3% isoflurane in a O_2_-enriched chamber and transcardially perfused with ice cold, oxygenated (95% O2/5% CO_2_) artificial cerebral spinal fluid (ACSF, 2-5°C) for 2 min at a rate of 5 ml/min prior to decapitation. ACSF was composed of (in mM): 126 NaCl, 2.5 KCl, 25 NaHCO_3_, 1.3 NaH_2_PO_4_, 2.5 CaCl_2_, 1.2 MgCl_2_, and 12 glucose. Mouse brains were then extracted, marked for laterality and sectioned in ice cold ACSF. Horizontal slices (300 μm thick) containing the thalamic ventrobasal complex (VB), including the ventrolateral nucleus, were cut with a Vibratome VT1200S (Leica) and transferred to a beaker containing warmed (35°C), oxygenated ACSF for 30 minutes and then at room temperature until transferred to a recording chamber.

### Whole-cell electrophysiology

After recovery, brain slices were transferred to the recording chamber and superfused with ACSF (22-25°C) at a flow rate of 2.5 ml/min. Recordings were obtained from thalamocortical neurons from the VL area visually identified with a microscope and an infrared CCD camera. Recording electrodes made of borosilicate glass had a resistance of 3-5 MΩ when filled with intracellular solution. Membranes of thalamocortical neurons were ruptured while at -65mV after achieving a >1GΩ seal. For some of the recordings, neurobiotin was used in the internal solution (0.5 mg/mL) for anatomical visualization within the VL.

### Current clamp (dual step protocol)

Evaluation of tonic firing consists of the count of action potentials following injection of positive current in increasing +25pA steps to induce depolarization (each depolarization step and plotted against the amount of positive current (pA) injected at that step). Rebound firing was quantified as the number of spikes per burst (No. of SPB) potentials, also referred to as low-threshold calcium spikes (LTCS) from T-type channel activation, following injection of negative current in decreasing -25pA steps to induce hyperpolarization. Each hyperpolarization step and plotted against the amount of negative current (pA) at that step. Each depolarization and hyperpolarization step had the same 500ms duration in each sweep with an inter-step duration of 500ms. The x and y axis in current clamp recordings show time in x axis (ms) and membrane potential in y axis (mV). The internal solution for these recordings contained (in mM): 125 K-gluconate, 8 NaCl, 1 MgCl2, 10 HEPES, 0.1 EGTA, and was adjusted to 7.25 -7.3 pH with KOH and to 275-285 mOsm with sucrose. Neurons were kept at -65mV by constant injection of positive or negative current. For experiments with HCN inhibitor, the dual step protocol was run prior to and after superfusion with ZD7288 (10μM).

### Parameter and passive membrane properties analysis

Tonic and rebound rheobases measure the minimum amount of positive or negative current to elicit an action potential. Results on rheobase report amount of current required to elicit the first observed action potential in tonic and rebound firing, respectively. Calculation of AHP was evaluated from action potentials spaced away from the initial T-type mediated burst observed in tonic firing, usually above 300ms. A resulting trace was obtained by Clampfit after averaging the first 2-3 action potentials that appear, usually between the first and fourth sweeps. A similar principle was used for the phase plot analysis, where selected action potentials were spaced away from the initial burst. Clampfit (version 10.7) was used to create an action potential waveform template to search for 3-4 representative action potentials. A resulting single trace of the average of these action potentials was used for the phase plot analysis. The resulting trace was transferred to the results window displaying the time stamp in ms (x values) at each membrane potential (mV) value reached by the action potential. Function cC = diff(cB) was used to evaluate mV/dt=diff(mV) and resulting values were then plotted against its corresponding membrane potential (mV).

### Voltage-clamp experiments (inactivation and recovery from inactivation protocols)

T-type Ca^2+^ currents were isolated with an internal solution containing tetramethylammonium (TMA) at a concentration of 5mM and QX-314 (5mM) adjusted to 7.2 pH with hydrofluoric acid, and 300 mOsm. Series resistance was compensated up to 70% compensation. T-type Ca^2+^ channel inactivation protocol consisted of holding the neuron at varying Vm (−120 to -50mV) for ∼3500ms and then stepping the neuron to -50mV for 500ms). Ca^2+^ current amplitude, current density, V_50_ (calculated with Boltzmann sigmoidal equation: Y=Bottom+(Top-Bottom)/(1+exp((V50-X)/Slope), and I/Imax were analyzed. A recovery from inactivation protocol was used to evaluate kinetics of recovery from inactivation. A pre-pulse and a test pulse were induced by holding the neuron at -90 and stepping to -50mV. Inter-pulse intervals with increasing exponential values were calculated with a Matlab function (from 50ms to 1200ms) for the duration of the recording and used consistently across recordings. Current amplitudes were measured, and full recovery was determined when test pulse over pre-pulse amplitude ratio values reached 90%. Values of tau were calculated individually and fitted to a one-phase decay equation (*y= (y0-plateau)*exp(− k*x) + plateau)*.

### Optogenetics

Whole-cell patch-clamp configuration was followed by a 20ms light stimulation (at 470nm, 0.5 mW) at 50 ms. Elicited inward sodium currents were observed following the 50ms light stimulation timestamp. Inward currents showed a latency of 3 to 7ms, post-stimulation, indicating to be the result of monosynaptic transmission. These experiments were verified by adding NBQX in the bath, eliminating the synaptic response. For stimulation of cerebellar fibers, neurons were kept at -65mV and synaptic response was determined by the presence of an ePSC (inward Na^+^ current). For stimulation of SNr fibers, synaptic response was evaluated by holding the neuron at -50mV to determine the presence of an iPSC (outward Cl^-^ current).

### Immunohistochemistry

After recordings, slices were fixed in 4%PFA for at least 1 hour, followed by two 10-minute washes with phosphate buffered saline (1XPBS) at RT. Normal donkey antiserum at a concentration of 5% in 0.3% Triton in 1xPBS was used for blocking for 1 hr. This was followed by Streptavidin-Alexa488 secondary antibody incubation (1:500) for 1 hr, followed by two 10-minute washes. Slices were then mounted and visualized on an epi-fluorescent microscope.

### Experimental Design and Statistical analysis

Each n represents an individual recording from a specific neuron type. Comparisons were made between neuron types at a specific membrane potential (i.e., -80, -85, -90, and -95mV) or current step (i.e., 150, 200, 250, 300, and 350pA). Analysis was performed with blinding to optogenetic response. Normal distribution was determined based on passing the *Kolgomorov-Smirnov* test. For data sets with a normal distribution, we reported the mean ± SEM. For not normal distributions, we reported data sets. For tonic and rebound firing in Figure 2, unpaired t-test comparisons were evaluated at different current steps or membrane potentials (see statistics table for corresponding figure). For parametric data, Welch’s correction test was used to account for standard deviation differences. For nonparametric data, Mann-Whitney test was used to evaluate ranks. Tonic and rebound firing in Figure 6 before and after ZD7288, were evaluated using paired t-test and Wilcoxon-matched pairs for nonparametric data sets. All p values <0.05 are considered significant.

## Results

### Heterogeneity in tonic and rebound firing based on thalamic afferents

We sought to investigate if there were any differences in intrinsic excitability among ventrolateral thalamic neurons receiving cerebellar axonal projections versus those that did not. Adult wild-type C57BL/6 mice were intracranially injected in the cerebellar nuclei (Fig.1A *left*) with AAV-hSyn-CHIEF-mRuby three weeks prior to performing whole-cell patch recordings. We recorded from neurons within the contralateral ventrolateral thalamus (VLT) using mRuby fluorescence as a landmark. Patched neurons were maintained at -65mV (Fig. 1A *right* and *1B*) and then subjected to optogenetic stimulation to evoke neurotransmitter release from cerebellar terminals expressing channel rhodopsin. The presence or absence of opto-genetically induced excitatory postsynaptic current (ePSC) was used to classify thalamocortical neurons with optogenetic as Cb_**aff**_+ for recipient of cerebellar afferents, and those with no response as Cb_**aff**_- (Fig. 1B). A subset of recordings included neurobiotin in the internal solution to map neurons within the VLT (Fig. 2C). We observed no clear anatomical delineation distinguishing neurons with cerebellar input from those without. In fact, these two populations (Cb_**aff**_+ and Cb_**aff**_-) could be found adjacent and intermingled with each other, suggesting a heterogeneous population within the VLT. Following optogenetic stimulation, thalamic neurons were maintained at -65mV prior to starting the dual-step protocol in current clamp (Fig. 2A).

**Figure 1.**
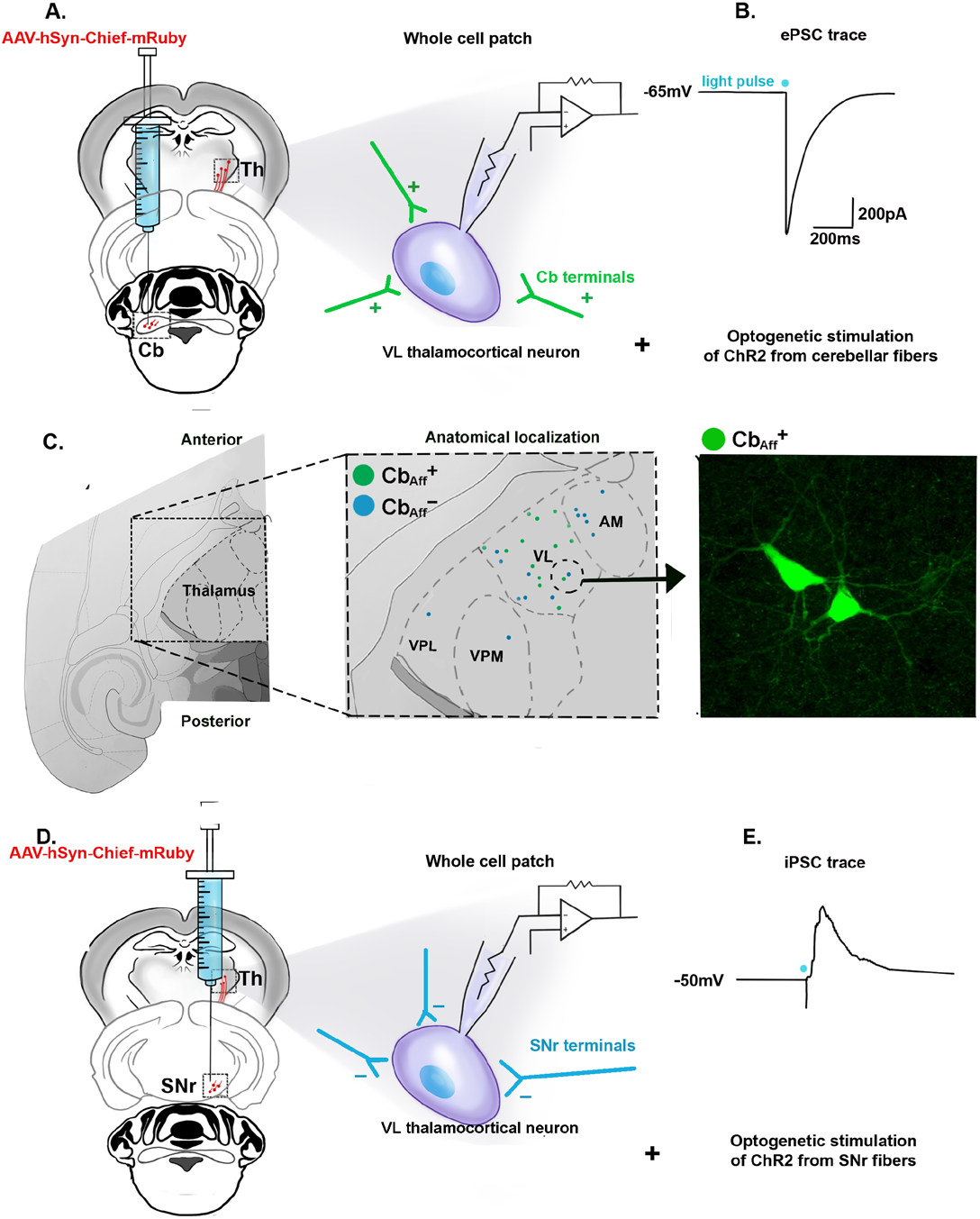
Whole cell patch clamp configuration followed by optogenetic stimulation of axonal fibers projecting from cerebellar or SNr nuclei. A) AAV-hSyn-CHIEF-mRuby was injected unilaterally in the lateral/interposed nuclei of the cerebellum. Patch clamp recordings were performed on thalamocortical neurons from the contralateral (to injection site) VL thalamus within the mRuby fluorescence area. B) To determine synaptic cerebellar input, each neuron was held at -65mV and tested for the presence or absence of an ePSC while eliciting a 470nm light pulse. C) Mapping of neurobiotin filled cells within VL thalamus demonstrates that cells with and without Cb afferents adjacent to one another. D) AAV-hSyn-ChR2-mCherry was injected unilaterally in the SNr nuclei of the basal ganglia and patch clamp recordings were performed on thalamocortical neurons from the ipsilateral (to injection site) VL thalamus within the mCherry fluorescence. E) Same as in B), except that each neuron was held at -50mV and tested for the presence or absence of an iPSC during light stimulation.

**Figure 2.**
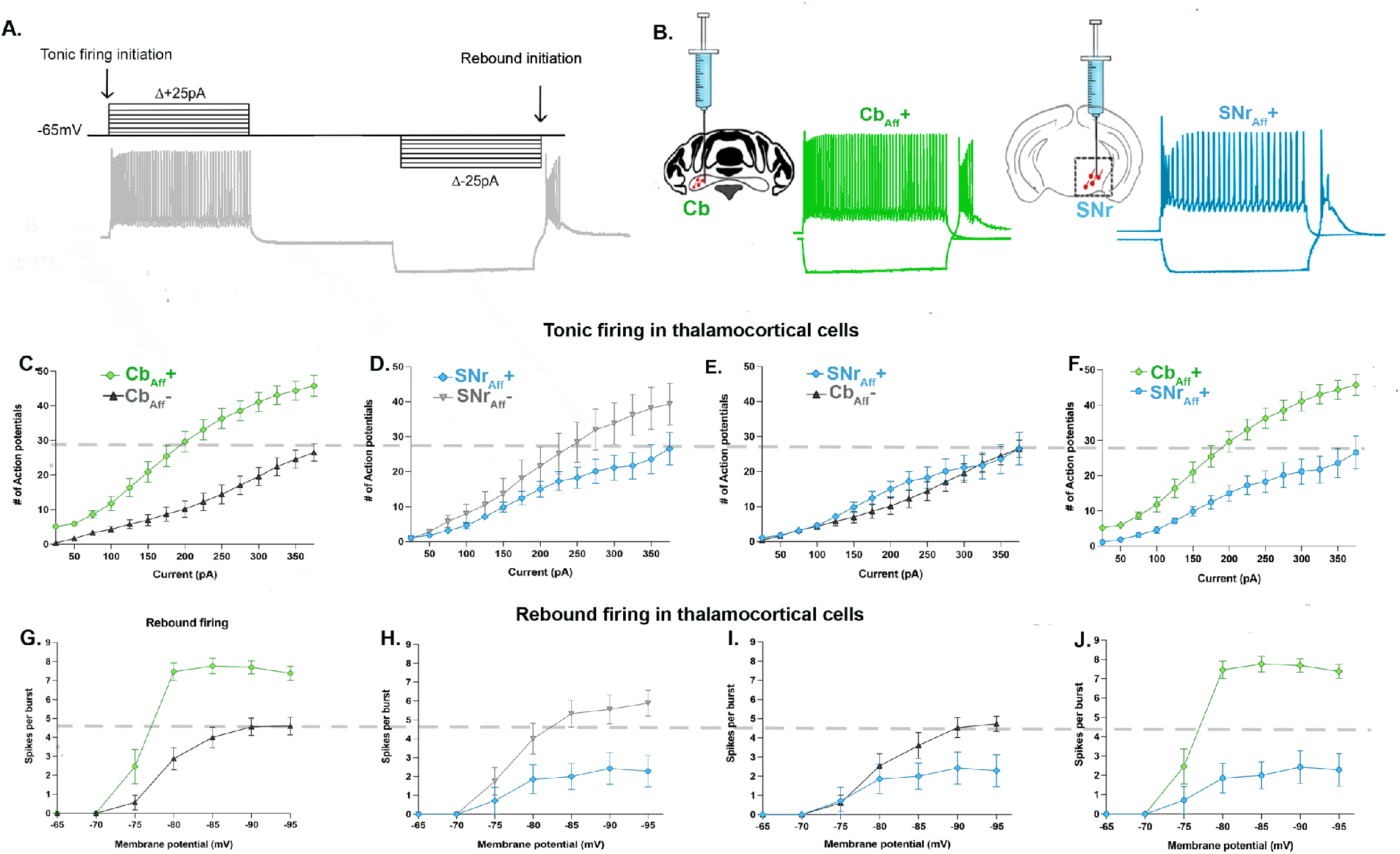
Intrinsic excitability of thalamocortical neurons with cerebellar and SNr input. A) Dual-step protocol in current clamp showing resulting tonic and burst firing modes. B) Representative traces of tonic and rebound firing from neurons with cerebellar (Cb_**aff**_+) or SNr input (R-SNrTh) input. C) Differences were observed in the No. of APs (***p<0.0001) between Cb_**aff**_+ and Cb_**aff**_-when compared at 150pA-350pA. D) Comparison between SNr_**aff**_+ and SNr_**aff**_- or E) SNr_**aff**_+ and Cb_**aff**_- showed no differences at any current step. F) Pronounced differences in the No. of APs were present when comparing Cb_**aff**_+ and SNr_**aff**_+ at 150pA-350pA. G) Significant differences were also found in the No. of SPB (***p<0.0001) displayed between Cb_**aff**_+ vs. Cb_**aff**_- at -80mV, -85mV, - 90mV and -95mV. H) Differences in SPB were detected at -85mV, -90mV and -95mV between SNr_**aff**_+ and SNr_**aff**_- I) Comparison of No. SPB between SNr_**aff**_+ and Cb_**aff**_- showed a difference only at -95mV (*p=0.0228). J) Differences in SPB were detected at - 80mV, -85mV, -90mV and -95mV when comparing only Cb_**aff**_+ and SNr_**aff**_+ neurons response. A bonferroni correction was used for multiple comparisons and a p value of <0.025 was considered significant for these data sets.

Tonic firing was measured by quantifying the number of action potentials (No. of APs) during the depolarization steps. Rebound burst firing was observed following hyperpolarization steps, where the number of spikes per burst (No. of SPB) were quantified. Analysis of tonic firing between Cb_a**ff**_+ and _Cba**ff**_- showed remarkable differences (Fig. 2C). Approximately, 60-160% more APs were observed during depolarizing steps in Cb_A**ff**_+ neurons when compared to Cb_A**ff**_-, with the largest differences at the smaller step sizes. Significant differences were found between groups (***p<0.0001), compared at +150pA through +350pA (see Table 1 for statistical comparisons for each step size). Interestingly, results also showed approximately 80-198% higher No. of SPB in the Cb_a**ff**_+ than Cb_**aff**_- between -80mV and -95mV (***p<0.0001), with the highest differences at -80mV (Fig. 2G) (Table 1).

**Table 1.**
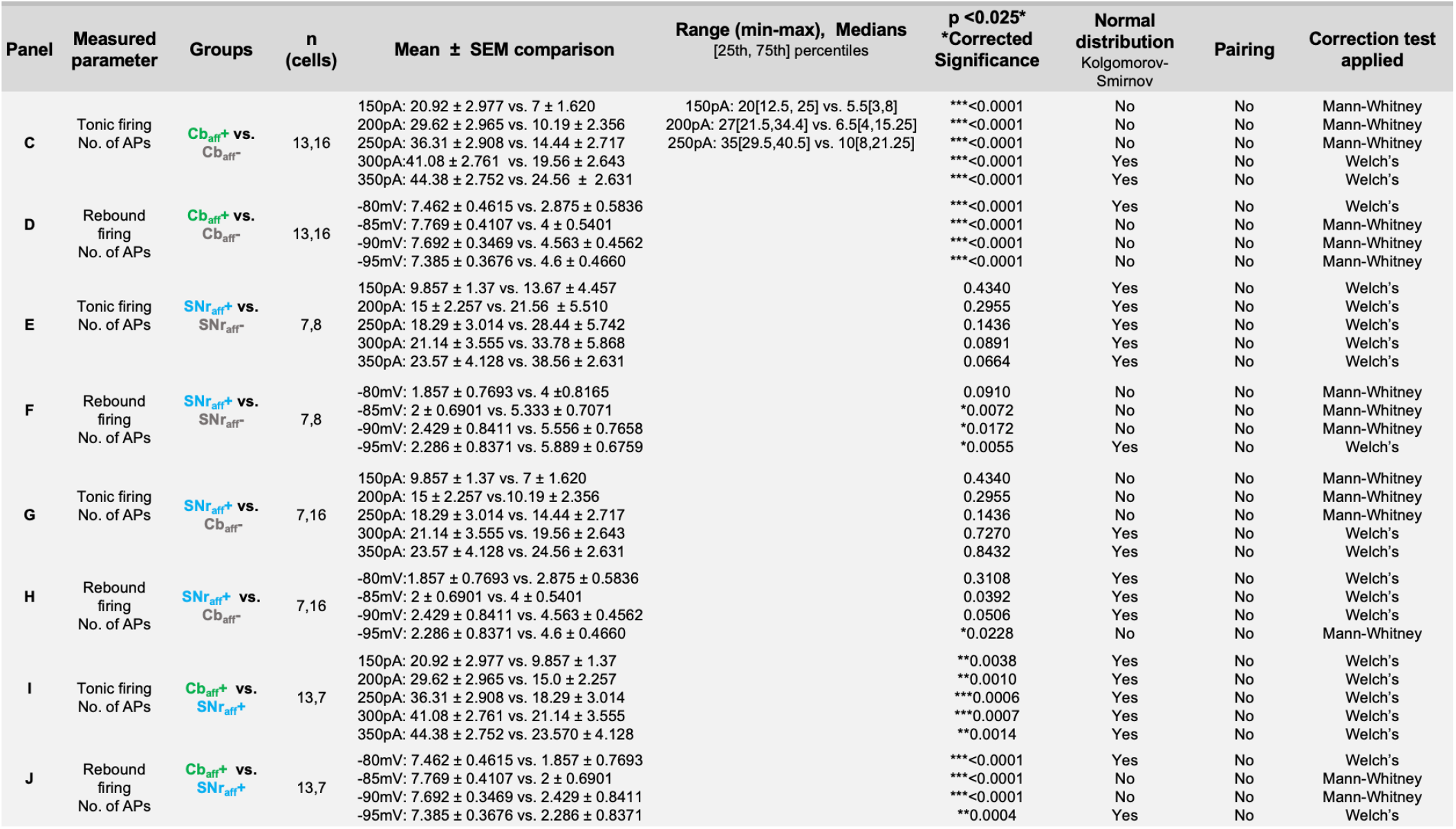
Statistical analysis for data in Fig. 2.

Multiple studies have reported the basal ganglia (SNr and GPi/EPN) as the other major input to the VLT (Kim *et al*., 2017; Pelzer *et al*., 2017; Hintzen *et al*., 2018). Therefore, we hypothesized that neurons without cerebellar input could be classified as basal ganglia-recipient neurons. To confirm input from the basal ganglia, we injected the same viral construct into the SNr nuclei (Fig. 1C and 1D) and the same optogenetic protocol was used to classify these thalamic neurons (SNr_**aff**_+ and SNr_**aff**_-). Neurons were patched within the VLT and SNr fibers were stimulated in the area (Fig. 1C *left*). For these experiments, neurons were classified as SNr_**aff**_+ for “response” if an iPSC was present following optogenetic stimulation (Fig. 1D). When comparing tonic firing in SNr_**aff**_+ and SNr_**aff**_- we observed no differences in tonic firing between the groups (Fig. 2D), however more rebound firing was observed in SNr_**aff**_- neurons between - 80mV and -95mV. We further compared Cb_**aff**_- and SNr_**aff**_+ neurons and saw no difference in No. of APs during tonic firing between them (see Table 1) (Fig. 2E). Rebound firing is significantly higher in the Cb_**aff**_- group, but only at -95mV (Fig. 2I). As we had predicted, Cb_**aff**_- and SNr_**aff**_+ showed similar intrinsic excitability levels, suggesting they may reflect the same population.

Our classification of thalamic afferents relies on a virus-mediated approach which may yield false negatives in our non-responder groups due to incomplete transfection of the afferent projections. To rule out the impact of false negatives when comparing groups, we directly compared tonic and rebound firing between Cb_**aff**_+ and SNr_**aff**_+ only, groups with opto-genetically confirmed cerebellar or SNr input. Interestingly, the No. of APs quantified in neurons with cerebellar input were still significantly higher than those displayed by neurons with SNr input, with depolarizing current injections between 150-350pA (Fig. 2F). Similarly, the No. of SPB were remarkably higher in the Cb_**aff**_+ than in the SNr_**aff**_+ neurons, with values of about ∼300% more at potentials between -80mV and -95mV (Fig. 2J). These results suggest at least two distinct populations of thalamic neurons whose intrinsic excitabilities can be attributed to their type of afferent. Our comparison analysis of neurons with presumed and confirmed afferent input, support the hypothesis that neurons in the Cb_**aff**_- are predominantly innervated by a region of the basal ganglia (i.e., SNr) and are a good representation of neurons with this type of input. To our knowledge, firing differences between these two populations within the VLT have not been previously investigated.

### Passive membrane properties

Current clamp recordings were further analyzed to compare the biophysical properties of neurons with cerebellar or basal ganglia input. We found that input resistance was higher (**p=0.0084)(Fig. 3A,B) and capacitance was lower (*p=0.01250)(Fig. 3C) in the Cb_**aff**_+ than in the Cb_**aff**_- neurons. No significant differences were found when comparing the resting membrane potential at I=0 pA (p=0.5150)(Fig. 3D).

**Figure 3.**
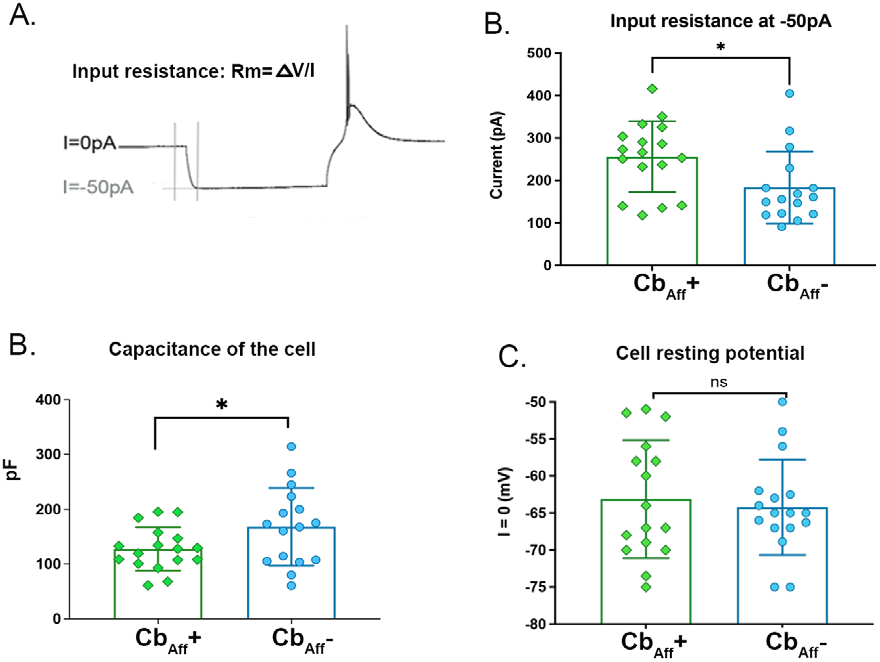
Passive Membrane properties of cells with and without cerebellar afferents. A) Example trace shows how input resistance was measured on -50pA hyperpolarization. B) Input resistance is higher in the Cbaff+ neurons. C) Cell capacitance was lower in Cbaff+ neurons. D) Cell resting potential was not different between populations.

**Figure 4.**
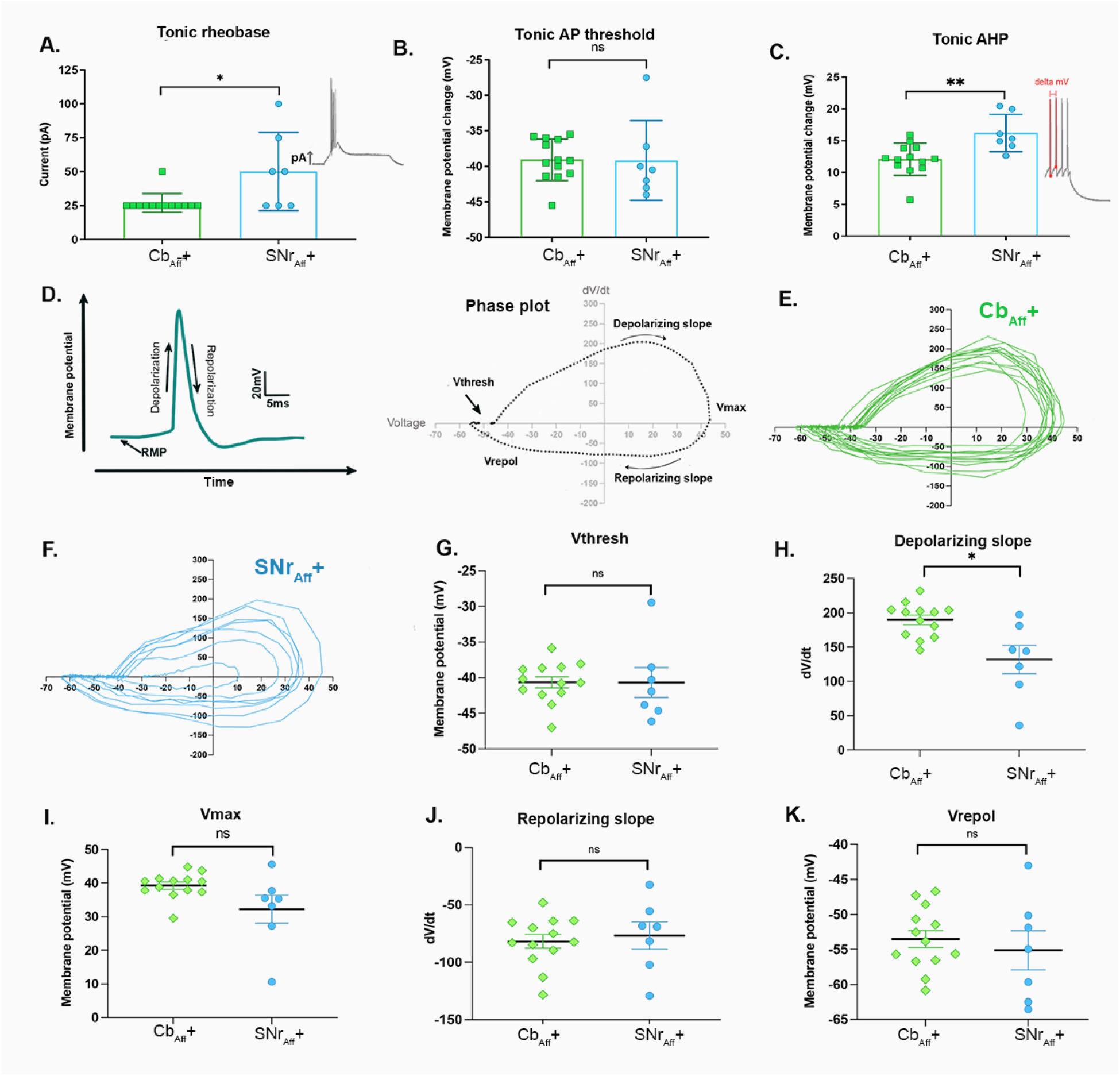
Phase plot analysis to evaluate active membrane properties in CbTh and SNrTh neurons. A) Significant differences in rheobase were observed (*p=0.0190). B) No significant differences were found in tonic AP threshold. C) Significant differences were found between groups when evaluating afterhyperpolarization (AHP) induced voltage changes during tonic firing. D) Left: graph showing the stages of an action potential over time (left). Right: representative phase plot analysis obtained by plotting membrane potential against its derivative over time, during an action potential. E) Individual phase plots for each cell with cerebellar input. F) Individual phase plots for each cell with SNr input. G) No significant differences were found in Vthresh. H) Significant differences were found in the depolarization slope (maximum dV/dt) between groups. I-K) No differences were found when comparing Vmax, the repolarizing slope (minimum dV/dt), or V_repol_.

### Action Potential Threshold and Waveform

We further analyzed tonic firing properties of Cb_**aff**_+ or SNr_**aff**_+ neurons to gain insight into ion channel mechanisms driving differences in tonic or rebound firing between groups. We first assessed two aspects of action potential initiation: rheobase and threshold. We observed a lower rheobase for tonic firing in the Cb_**aff**_+ neurons, indicating that less current was required to initiate tonic firing (*p=0.019) (Fig. 4A). These results correlate with higher input resistance observed in the Cb_**aff**_+ group (Fig. 2B). No differences were found in threshold potential for firing (p=0.96) (Fig. 4B). Interestingly, we found that afterhyperpolarization (AHP) was significantly smaller in Cb_**aff**_+ neurons, compared to SNr_**aff**_+ neurons (**p=0.0089) (Fig. 4C). A phase plot analysis of the action potential waveform during tonic firing (Fig. 4D-F) was analyzed for every neuron in each group. We found that Cb_**aff**_+ neurons exhibited a steeper depolarization slope that was significant when compared to SNr_**aff**_+ neurons (*p=0.0301). (Fig. 4H). In contrast, quantification of V_thresh_, V_max_, repolarizing slope, and V_repol_ (Fig. 4G, I-K) revealed no significant differences between neuronal types. The observed differences in action potential AHP and depolarization slope strongly suggest possible differences in Na^+^ and K^+^ channel expression or function between neurons with cerebellar or basal ganglia inputs.

We also performed analysis of the rebound burst firing after hyperpolarizing steps. Similar to tonic firing, we observed a significantly lower rebound rheobase (Fig. 5A) in the Cb_**aff**_+ group (p=0.0028), but no differences in the action potential (AP) threshold (p=0.0693) (Fig. 5B). Furthermore, we quantified the latency of the first burst to appear following hyperpolarization. While the same amount of current in Cb_**aff**_+ and SNr_**aff**_+ elicits higher rebound firing in Cb_**aff**_+ neurons, the latency for each burst is significantly delayed in the Cb_**aff**_+ group (p=0.0082)(Fig. 5C).

**Figure 5.**
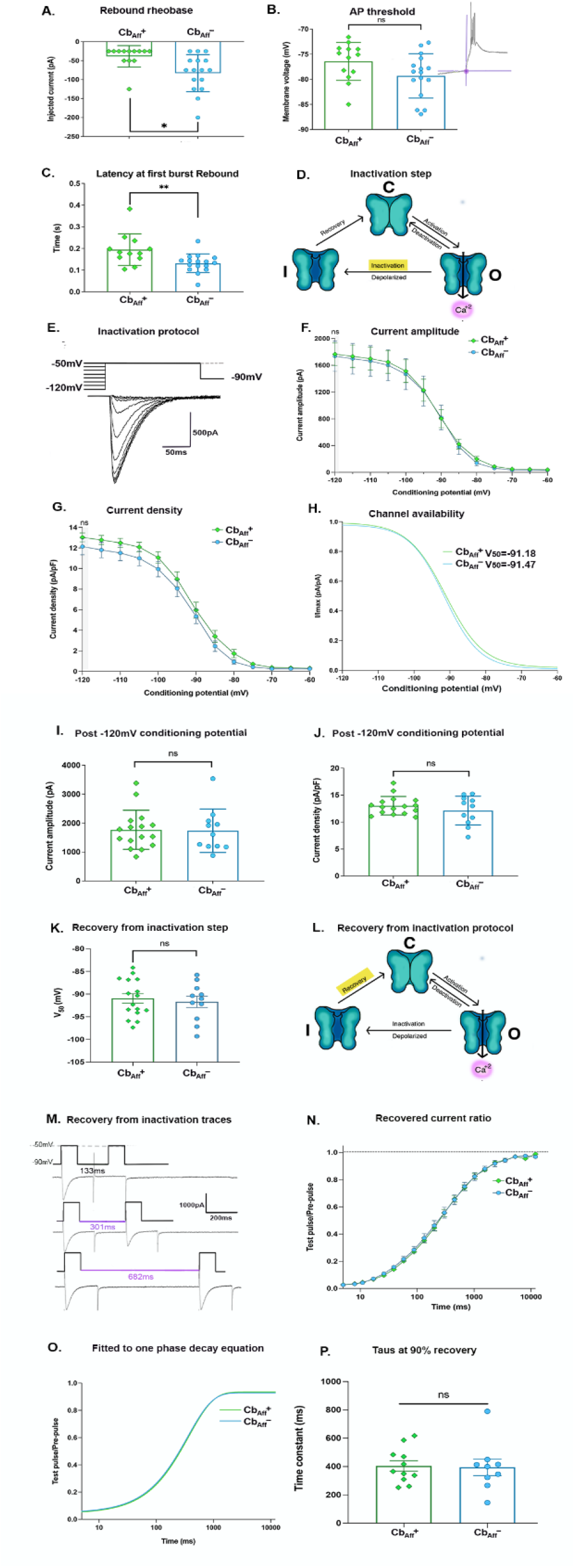
Rebound firing and T-type channel mediated Ca2+ current amplitude and density analysis. A) Cbaff+ neurons require smaller current injections to induce rebound burst firing than Cbaff-neurons. B) AP threshold determined membrane potential measurement prior to first rebound burst or at least one action potential was not different between groups C) Latency to first burst was faster in Cbaff-compared to Cbaff+ neurons. D) T-type Ca^2+^ ion channel schematic showing its closed, open, and inactivated states, the “inactivation” stage emphasized in red. E) Voltage clamp protocol (top) showing various conditioning potentials ranging from (−120mV to -50mV) and test potential at -50mV (500ms). Inter-sweep membrane voltage was maintained at -90mV. Representative traces (bottom) of Ca^2+^ current amplitudes after various conditioning potentials F) Current amplitudes plotted against conditioning potential were not different. G) Current density (pA/pF) were also not different. H) Channel availability or I/Imax ratios were graphed and fitted to the Boltzmann sigmoidal equation to calculate average V_50_ values per group V_50._ Individual calculated values showed no differences I) maximum current and J) current density at -120mV was not different between groups. K) V50 for inactivation was not different between groups. L) T-type Ca^2+^ ion channel schematic showing its closed, open, and inactivated states, the “recovery from inactivation” stage emphasized in yellow. M) Recovery from inactivation protocol also in voltage clamp showing a pre-pulse and a test pulse at -50mV and representative traces of Ca^2+^ current amplitudes at the pre-pulse and test pulse, until reaching at least 90% recovery. N) Ratios of test pulse/pre-pulse between groups (between 0 to 1) and O) were fitted to one phase decay to calculate tau P) Individual taus at 90% recovery were calculated per group, no significant differences were found.

**Figure 6.**
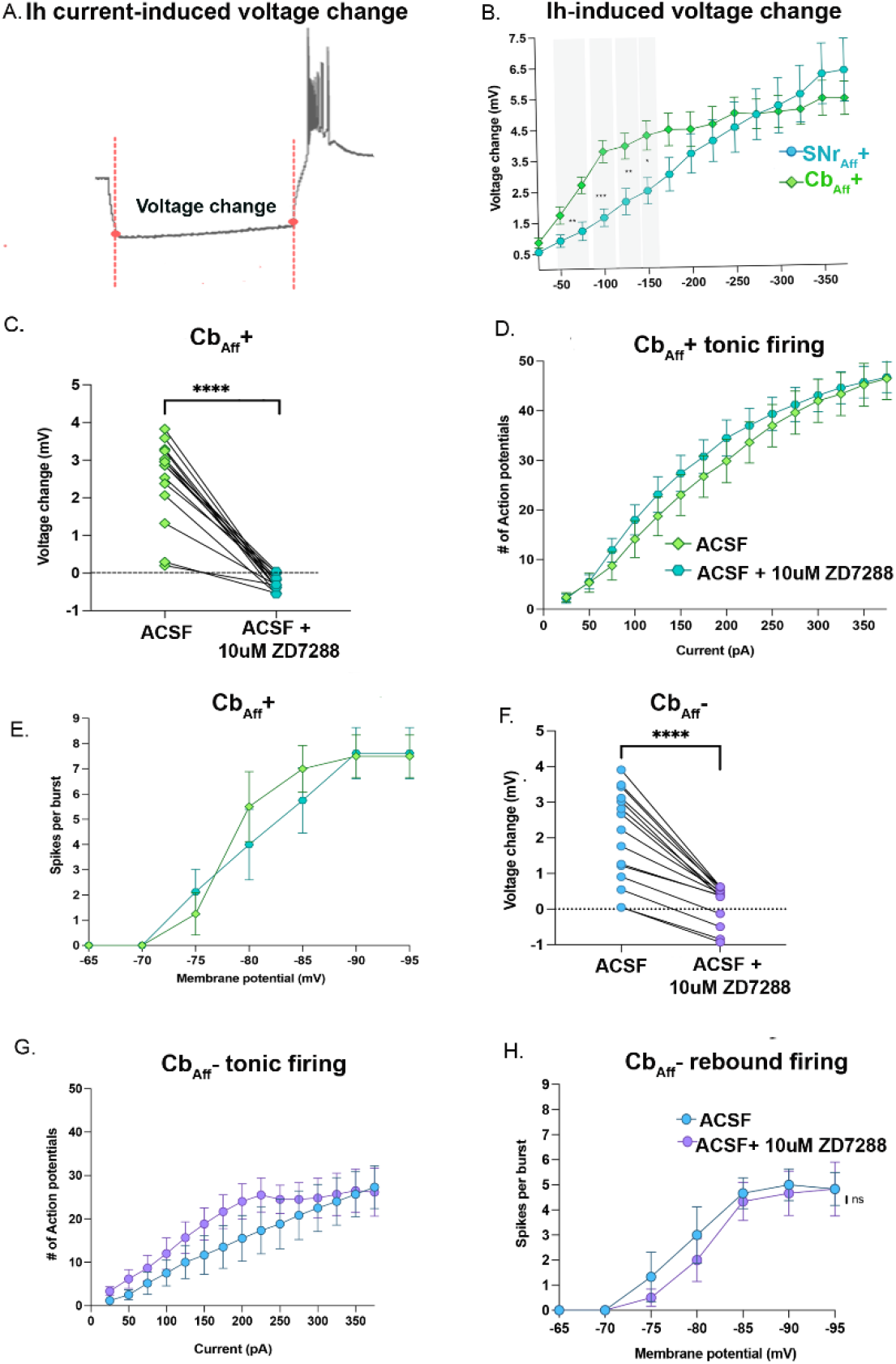
Intrinsic excitability measured before and after inhibiting HCN channels with ZD7288. A) Representative trace from a current clamp recording showing a membrane potential difference at the end of the hyperpolarizing or negative current step. B) Significant differences in sag amplitudes were observed between -50pA and -150pA. C) ZD7288 was tested for its efficacy to reduce hyperpolarization sag in Cbaff+ neurons. D) ZD7288 had no effect on tonic or E) rebound burst firing in Cbaff+ neurons F) ZD7288 was tested for its efficacy on Cbaff-neurons G) ZD7288 had no effect on tonic or H) rebound burst firing in Cbaff-neurons

### Kinetic properties of T-type Ca^2+^ currents are not different between groups

The dramatic difference in rebound firing between our populations led us to hypothesize that T-type Ca^2+^ channels activity may be greater in neurons with cerebellar afferents. To test this, we performed experiments in voltage-clamp to isolate T-type-mediated Ca^2+^ current amplitude and current density between groups. We used a voltage-clamp T-type channel inactivation protocol to measure Ca^2+^ currents in Cb_**aff**_+ or Cb_**aff**_- neurons, distinguished with the same optogenetic methodology as in Fig. 2A. The voltage clamp inactivation protocol holds thalamic neurons at progressively increasing hyperpolarization conditioning potentials prior to stepping the membrane potential up to -50mV (testing potential), at which point, peak Ca^2+^ current amplitudes are measured (Fig. 5E). Resulting current amplitudes in each cell were plotted against conditioning potential, but no differences were found in current amplitude (p=0.9161) (Fig. 5F, I) or current density (p=0.3444) (Fig. 5G, J) between groups. Channel availability (I/Imax) curves were fitted to Boltzmann sigmoidal equation to calculate overall V_50_ mean values for each neuron per group (Cb_**aff**_+: -91.18 mV and Cb_**aff**_-: -91.47mV) (Fig. 5H). Analysis of individual V_50_ values showed no significant differences (p=0.6529) (Fig 5K).

Differences in T-type isoform expression may not be detected with I_max_ or V_50_ measurements. There are three types of low-voltage activated (LVA) T-type Ca^2+^ channel isoforms that express abundantly in the thalamus, Ca_V_3.1, Ca_V_3.2, Ca_V_3.3. Ca_V_3.1 and Ca_V_3.2 isoforms have similar recovery from inactivation times, whereas Ca_V_3.3 are three times slower. While Ca_V_3.1 is most highly expressed in the ventrobasal thalamus (where VL is located) it is possible that a difference in the relative expression of Ca_V_3.1 and Ca_V_3.3 between neuronal populations, could drive rebound firing differences. Therefore, to explore differences in isoform expression, we measured recovery from inactivation times. Neurons were first maintained at -90mV and then depolarized to -50mV to obtain an initial inward Ca^2+^ current (pre-pulse)(Fig. 5M). This was followed by a hyperpolarization back to -90mV and then a depolarizing step to -50mV for a second inward Ca^2+^ current (test-pulse). Inter-pulse durations between pre and test-pulses was exponentially increased per sweep. Ratios of recovered currents were plotted against time (Fig. 5N) and fitted to a one-phase decay equation (*y= (y0-plateau)*exp(−k*x) + plateau)* when reaching at least a 90% of current recovery (Fig. 5O). Recovery time (tau) was calculated for each cell, but no significant differences were found between groups (p=0.8837)(Fig. 5P). Altogether, these data suggest that differential expression of T-type channels is not likely driving the increased levels of rebound firing observed in the Cb_**aff**_+ population.

### HCN channels in intrinsic excitability

Another family of ion channels that promotes rebound firing in the thalamus is the hyperpolarization-activated cyclic nucleotide gated (HCN/I_h_) channel (Biel *et al*., 2009). When activated by membrane hyperpolarization, these cation channels open to restore the membrane potential, appearing as a sag during sustained hyperpolarization. To evaluate HCN activation during hyperpolarization steps, we compared membrane voltage values at the beginning and at the end of each hyperpolarization step (Fig. 6A). We observed significant differences in the amount of voltage sag, evaluated at the onset and end of the initial hyperpolarization steps, between Cb_**aff**_+ and SNr_**aff**_+ neurons (Fig. 6B). These differences were apparent at -50pA (**p=0.0084), -75pA (**p=0.0045), -100pA (***p=0.0003), -125pA (**p=0.0085), and at -150pA (*p=0.011) (Fig. 6B). We next hypothesized that HCN channel inhibition would reduce rebound firing or the No. of SPB in the Cb_**aff**_+, with number of spikes comparable to SNr_**aff**_+ group. To test this, we applied ZD7288 at a concentration of 10uM (a selective HCN inhibitor) in the bath and verified HCN inhibition by measuring voltage sags before and after bath application of the inhibitor. ZD7288 effectively inhibited HCN-mediated Ih currents and significant differences were observed as a result of HCN channel inhibition in the Cb_**aff**_+ and Cb_**aff**_- (****p<0.0001) (Fig. 6C,F). However, no effects were observed in neither tonic nor rebound firing and excitability differences between Cb_**aff**_+ and Cb_**aff**_- remained (Fig. 6D, E, G, H). Overall, we conclude that the HCN channels activation do not seem to play a role in mediating rebound firing differences between two groups.

## Discussion

We report that thalamocortical neurons in the VL with cerebellar input display higher gain (slope of current-frequency relationship) and maximum tonic and rebound firing rates than thalamocortical cells with basal ganglia input. Our results suggest the existence of a heterogenous population of thalamocortical neurons within the mouse VL thalamus that can be distinguished by their firing properties and afferent inputs (i.e. cerebellum vs. SNr). Pathway specific single-cell RNAseq analysis revealed cells with Cb vs SNr inputs (primarily localized to VL and VA, respectively) had differential expression of several voltage-gated ion channels that contribute to firing properties (Phillips *et al*., 2019). Our electrophysiological findings support that gene expression differences observed in these thalamic populations have functional impacts on excitability. Finally, higher input resistance allows for a greater shift in Vm for any given current injection (V=IR), which likely contributes to lower rheobase for tonic and rebound firing and larger gain in current-frequency relationships in Cb_**aff**_+ neurons.

### Higher excitability associated with cerebellar afferents

Input from the cerebellum and the basal ganglia innervating the VL was expected; however, we did not predict the heterogeneity or differences in intrinsic excitability between adjacent cells. Neurons as close as ∼30μm apart had different input and excitability (Fig. 1C). In correlation with our findings, an *in vivo* study in macaca monkeys showed that groups of thalamic neurons with cerebellar afferents were more excitable than those with GPi input (basal ganglia). In fact, this group classified these areas as ‘microexcitable’ zones within the caudal part of the VL thalamus (Buford *et al*., 1996) and ruled out pallidal input to these areas. They found that current stimulation of these microexcitable zones required only a third of the amount of current that pallidal receiving areas did to elicit the muscle contraction or twitch in the monkeys’ arms. Afferents to these areas with higher excitability were traced back to the cerebellar dentate nuclei. Our data also shows lower rheobases in both tonic and rebound firing rates in Cb_**aff**_+ neurons, when compared to SNr_**aff**_+ neurons, consistent with findings in primates (Buford *et al*., 1996). Furthermore, our analysis of the passive membrane properties suggests that lower rheobases may be attributable to a higher input resistance and a smaller cell capacitance, allowing for greater changes in membrane potential at any given amount of current (Ohms law) in Cb_**aff**_+ neurons.

### Cerebellar and basal ganglia input distribution

Thalamic excitability is modulated by a combination of excitatory and inhibitory input from different areas. Excitatory afferents primarily project from the cortex and the cerebellum, and inhibitory afferents derive from the basal ganglia (GPi and SNr) (Garcia-Munoz & Arbuthnott, 2015; Kim *et al*., 2017; Pelzer *et al*., 2017). It is well known that cerebello-thalamo-cortical and basal ganglio-thalamo-cortical projections converge within the somatosensory and motor cortices. However, it is far less understood if and how both projection systems interact at the level of the thalamus. In fact, anatomical and electrophysiological studies have shown interspecies (non-human primates, cats and some in dogs) discrepancies on whether cerebello-thalamic and basal ganglio-thalamic projections synapse in segregated areas within the thalamus or if they converge onto the same TC neurons (Hintzen *et al*., 2018). Similarly, studies in rats have also shown evidence supporting either convergence or segregation (Chevalier & Deniau, 1982; Kuramoto *et al*., 2011; Hintzen *et al*., 2018). Anterograde labeling from SNr and cerebellar dentate nuclei showed slight overlap of these projections at the interface between VL and VA (Phillips *et al*., 2019). Our electrophysiological data also suggests that even at this interface, there are distinct firing properties between confirmed populations with cerebellar input or SNr input. Our findings with increased excitability of Cb_**aff**_+ neurons are consistent with previous characterization across thalamic nuclei demonstrating that density of innervation from cerebellar afferents in VL was associated with increased excitability relative to medial thalamic nuclei with fewer cerebellar afferents (Gornati *et al*., 2018). The previous study did not examine excitability of neurons within the VL that do not receive cerebellar afferents. Our data suggest that even within the VL, cerebellar innervation is associated with higher excitability.

While tonic firing between SNr_**aff**_+ and Cb_**aff**_- were similar, rebound firing rates were different between these populations. The SNr is located in a much broader area than the cerebellar nuclei and we speculate that the viral construct did not transfect the same proportion of neurons in each area. Therefore, our experimental design in the SNr could be more prone to false negatives, likely skewing our results, while Cb_**aff**_- seemed provided a good representation of neurons with basal ganglia input. While we have identified at least 2 populations of VL thalamic neurons based on afferent inputs and firing properties, we cannot rule out the possibility that some neurons in this area receive input from both sources. TC neurons in the VL also receive afferents from the entopeduncular nucleus (EPN), human GPi alternative in the mouse, therefore, we cannot conclude that every neuron in the Cb_**aff**_- receives only SNr input. Further studies using anatomical tracers and single cell electrophysiology are needed to provide further clarity regarding the convergence or segregation of inputs from cerebellum and basal ganglia in the thalamus.

### Action potential initiation and waveform

Voltage-dependent sodium and potassium channels are the main molecular mediators of action potential firing and neuronal intrinsic excitability. Phase plot analysis provides insight into sodium and potassium channel mechanisms influencing firing differences. Interestingly, our analysis showed that Cb_**aff**_+ cells depolarized faster, showing higher dV/dt values than SNr_**aff**_+ cells. Voltage slope values were measured at the rising phase of the action potential once the reached threshold. Because changes in membrane potential during the depolarization stage are mediated primarily by Na^+^ channels, our results strongly suggest the possibility of higher expression of Na+ channels or a difference in Na+ channel kinetics (or isoforms). Modeling of cortical neuron firing suggests that the depolarization slope is dependent on sodium channel cooperativity, a feature that is dependent on the density of this channel (Naundorf *et al*., 2006). Therefore, slope differences may reflect differences in channel density rather than expression. Kv1.2 is strongly expressed in thalamocortical neurons (Wang, 1994) and pharmacological inhibition of Kv1.2 regulates rheobase current and firing rates without altering threshold (Kasten *et al*., 2007). We observed differences in rheobase but similar action potential thresholds, consistent with a possible difference in Kv1.2 expression. We also observed a smaller after-hyperpolarization (AHP) voltage observed in Cb_**aff**_+ neurons. AHPs are largely mediated by small conductance calcium-activated potassium channels (SK) (Maylie *et al*., 2004). Application of apamin to thalamocortical neurons reduces the AHP and increases gain and maximum tonic firing rates (Kasten *et al*., 2007). We therefore speculate that the smaller AHP in Cb_**aff**_+ neurons likely contribute to the heightened excitability in this population. Differences in the AHP will also have a large influence on rebound burst firing which produces a T-type channel mediated increase in calcium that activates SK and can limit burst firing (Wolfart & Roeper, 2002; Hallworth *et al*., 2003). Together, our results suggest that the functional balance of sodium and potassium channels in Cb_**aff**_+ neurons are shifted to a more excitable state than SNr_**aff**_+ neurons.

### T-type and HCN channels in excitability

Measured T-currents following inactivation protocols, showed no significant differences in current amplitude, current density, I/Imax, or V_50_ between populations. This was surprising, given the dramatic differences in rebound or burst firing between groups and scRNAseq showing differences in CaV3.3 gene expression (Phillips *et al*., 2019). Consistent with our data, *in vivo* single cell recordings performed in VL neurons with basal ganglia input from the medial globus pallidus-GPm) revealed no significant differences in rebound firing rates between thalamocortical cells from WT and Ca_V_3.1 KO mice. Rather, they found differences in the latency of the rebound firing onset following synaptic inhibition (Kim *et al*., 2017). Interestingly, we also found longer latencies to the onset of rebound firing in Cb_**aff**_+ when compared to SNr_**aff**_+ cells following hyperpolarization. These results suggest that differences in rebound firing, observed between our Cb_**aff**_+ and SNr_**aff**_+, are independent of T-type channel expression. Similarly, while the HCN-mediated sag during hyperpolarization was significantly larger in Cb_**aff**_+ neurons, the HCN inhibitor did not strongly decrease rebound firing rates and still were dissimilar to SNr_**aff**_+ cells. The lack of effect by the HNC inhibitor is, perhaps, not surprising since the amplitude of the HCN sag was relatively small compared to other neuronal types where HCN makes a strong contribution to rebound or spontaneous firing.

Altogether, the differences in excitability observed between these populations likely influence synaptic integration and transmission of information to cortical areas. These differences could be primarily attributed to differences in the expression of sodium and potassium ion-channels and not T-type or HCN channels. Further electrophysiological studies are still required to find better correlations with differences in ion-channel expression profiles. It also remains to be elucidated how these populations may respond to neurological disorders influencing their afferent nuclei.

## Acknowledgements

We thank the Aoto Lab for providing us with the viral construct we used for optogenetic experiments. We thank Bryan C. Swanton for his contribution to our statistical analysis.

